# Pinching the cortex of live cells reveals thickness instabilities caused by Myosin II motors

**DOI:** 10.1101/2020.09.28.316729

**Authors:** V. Laplaud, N. Levernier, J. Pineau, M. San Roman, L. Barbier, P. J. Saez, A. M. Lennon, P. Vargas, K. Kruse, O. du Roure, M. Piel, J. Heuvingh

## Abstract

The cell cortex is a contractile actin meshwork, which determines cell shape and is essential for cell mechanics, migration and division. Because the cortical thickness is below optical resolution, it has been generally considered as a thin uniform two-dimensional layer. Using two mutually attracted magnetic beads, one inside the cell and the other in the extracellular medium, we pinch the cortex of dendritic cells and provide an accurate and time resolved measure of its thickness. Our observations draw a new picture of the cell cortex as a highly dynamic layer, harboring large fluctuations in its third dimension due to actomyosin contractility. We propose that the cortex dynamics might be responsible for the fast shape changing capacity of highly contractile cells that use amoeboid-like migration.

## Introduction

Dynamic cytoskeletal networks associated with the cell surface define the shape of mammalian cells ^1,2^. In particular, the actin cortex, a thin network of actin filaments just beneath the plasma membrane, plays a central role in shaping the cell surface ^3^, and in defining its mechanical properties ^4,5^. The actin cortex comprises, in addition to actin filaments, motors, membrane and actin linker proteins, actin nucleators ^6^ and cross-linkers, and regulatory proteins ^7^, which together render animal cell shape highly dynamic and able to adapt to external stimuli in a variety of physiological contexts such as cell migration or tissue morphogenesis.

Despite its central importance in cellular morphogenesis, the actin cortex remains poorly characterized, especially in terms of physical properties ^8^. Its physical dimension (thickness) was only recently measured in cultured mammalian cells, using optical methods ^9–11^, but whether and to which degree this thickness varies in time is not known. So far, cell cortex mechanics has been probed through shallow indentation of the cell with an Atomic Force Microscope ^12^ or through the twisting of ferromagnetic beads attached to the cell surface ^13^, but it is difficult in such experiments to separate the contribution of the cell cortex from the one of the rest of the cytoskeleton and cell organelles.

Since its discovery, the cortex has been mostly considered as a uniform two-dimensional structure ^4,8^. This is at least partly due to experimental limitations in imaging a structure whose thickness is smaller than the optical resolution, but also to the fact that contact probing can only be realized from the outside of the cell. In this work, we circumvent this obstacle by using a pair of probes (magnetic beads), one located inside the cell, and the other on the outside. The attraction between the beads is controlled by an external magnetic field, allowing us to slightly pinch the cortical layer. We can in this manner measure, with unprecedented spatial accuracy and temporal resolution, the thickness and dynamics of the cell cortex.

## Results

### The magnetic pincher: a robust method to measure physical properties of the cell cortex in live cells

Inspired by our previous work, in which we probed thin layers of actin networks assembled in vitro between superparamagnetic beads ^14,15^, we develop a new experimental set-up in which two magnetic beads pinch the cell cortex, one being inside the cell and the other outside (Fig. 1A, beads 1-2). In the present work, we probe the cortex of primary bone marrow-derived dendritic cells from mice. These cells display amoeboid motility: a migration mode independent of focal adhesions, stress fibers or large lamellipodial protrusions, which mostly relies on the fast remodeling of their actin cortex ^16–19^. Dendritic cells can also ingest large quantities of extra-cellular material including fluid (macropinocytosis) and particles (phagocytosis). This environment sampling activity allows them taking up antigens, which is the basis of their immune function, and in the present case enables entry of large magnetic beads, independently of specific receptor engagement. Cells loaded with uptaken beads and mixed with freely floating beads are placed in an external homogenous magnetic field. The field induces a tunable attractive force between the beads, which then pinch the cell cortex between them (Fig. 1B). The experiment is facilitated by the spontaneous organization of magnetic beads into pairs or chains when exposed to a magnetic field. Electron microscopy (TEM) imaging and fluorescent labelling confirm that ingested beads are surrounded by an endomembrane that has not fused with lysosomes (Fig. S1A-C), and that the cortex is indeed pinched between the internal and external beads (Fig. 1B).

**Figure 1.**
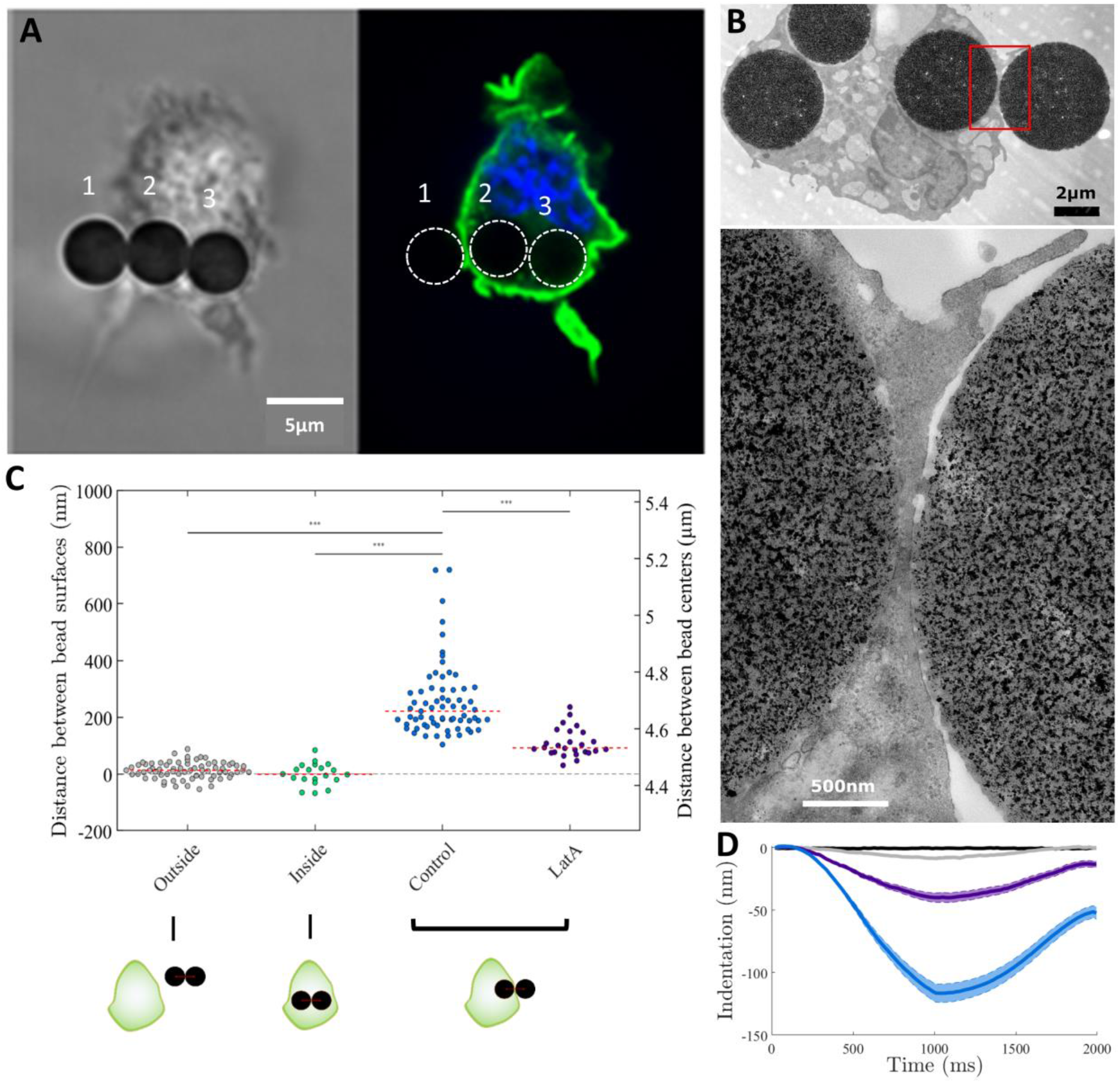
Measurement of cortex thickness using magnetic beads. A – Bright field (left) and fluorescence (right) images of a Dentritic Cell expressing Lifeact-GFP (green) and stained with Hoechst (blue), with internalized magnetic beads, under a magnetic field. Three beads, two in the cell (2 and 3) and one outside (1), are aligned by the magnetic field. The fluorescence image shows the pinching of the actin between beads 1 and 2 (white dotted outlines). Scale bar 5 μm. B – Transmission electron micrograph of the cortex of a cell pinched between two magnetic beads. The upper image shows the whole cell and the lower one is an enlargement of the red area. C – Distances between beads surfaces (left axis) computed from the measured distances between beads centers (right axis), for different configurations (schematized below the graph) and conditions. Each point represents the median of a 5 to 15 minute measurement at 1.25 Hz on a single cell. The red dotted line represents the median of the single cell measurements. Control cells (blue, number of cells n=67, number of experiments N = 10) display a median bead separation of 221 nm, whereas pairs of beads observed outside (grey, n = 69, N = 2) and inside (green, n =20, N =13) cells are in contact (average separation below the resolution). Cells treated with 500nM Latrunculin A (purple, n = 26, N = 5) show a large decrease in bead separation compared to control cells. D – Average of indentation curves obtained during compression experiments, in which the magnetic field is increased up to 50mT (~1000pN) in one second and decreased back to 5mT (70 pN) in one second. Bare beads (black, n_c_=49 compressions, n=16 bead pairs) are non-deformable (indentation below resolution). Serum coated beads outside of cells (beige, nc=143, n=11) show a nanometric deformable layer (max deformation below 10 nm), while beads on each side of the cortex (control, blue, nc=234, n= 20 cells, N = 5) display a reversible (elastic) deformation of 100 nm amplitude. 500 nM Lat A treatment strongly reduces the amplitude of the deformation (LatA, purple, n_c_=85, n=7, N = 1), down to about 40 nm.

A first important point is to determine the accuracy of the measured distance between the beads. The beads are monitored in 3D at a frequency of 1.25 Hz over typically 5 to 15 min with transmitted light. The measurement accuracy on the distance between the center of two beads is approximately 2 nm ^20^ in the plane of observation and around 45 nm in the perpendicular direction (see Sup Mat), resulting in an accuracy of 7 nm in the distance between bead centers. Subtraction of the bead diameter from the distance between the two centers gives the distance between the bead surfaces and thus the thickness of the pinched layer. We measure the beads to be highly monodispersed in size (Fig. S1D), thus ensuring the precision of the distance between the bead surfaces. Overall, our measurement accuracy on the absolute thickness of the pinched layer is 31 nm (see Sup Mat), which is way below the estimated thickness of the cell cortex. In addition, the thickness variation in time can be determined with a much better accuracy (7 nm, as the uncertainty on the bead diameters does not enter into this calculation) at >1 Hz, more precisely than any method used so far to estimate cortical thickness.

### Dendritic cell cortical layer has a median thickness of 220 nm

A small magnetic field of 5 mT is applied to produce a constant attractive force (~70 pN) between the beads. This force holds the beads in contact with the outer membrane and the inner face of the cell cortex so that the distance between the surfaces accurately reflects the cortex thickness. To estimate this thickness and control for potential artifacts, we perform three types of experiments:

1. We compare measurements of distances between two beads outside the cell, two beads inside the cell, and two beads pinching the cortex (Fig. 1C). While the distance between the surfaces of two beads inside or outside the cell are on average undistinguishable from zero, the distance measured for the cortical layer has a median value of 220 nm (Fig.1C, blue). This measurement is consistent with measurements in Hela cells with subpixel-resolution fluorescence imaging ^9^.
2. We increase the magnetic field, and thus the attraction force (up to 1nN), to check whether the structure pinched between the beads is deformable (Fig. 1D). Increasing the force between beads gives relative measures, which are precise down to a few nanometers. This shows that bare beads are stiff and non-deformable (less than 1 nm indentation), and beads used in the live-cell experiment show a minimal indentation (below 10 nm) due to the coating layer of serum (Fig. 1D, grey). In contrast, the cortical layer between a bead inside and a bead outside the cell is indented by more than 100 nm (Fig. 1D, blue), highlighting that a deformable object is being pinched. Upon force release, the distance between the beads relaxed, revealing the elastic nature of the compressed material.
3. We compare control cells and cells in which the actin cortex has been disassembled using a high dose of Latrunculin A (LatA) (Fig. 1C-D). Treatment with 500 nM LatA disassemble the actin cortex, as judged by fluorescent imaging (Fig S1E-F). The remaining layer, measured at 92 nm (Fig 1C, purple), which is significantly thinner than in untreated cells, is thicker than the distance between two beads inside the cell, and can be reversibly indented by about 40 nm (Fig. 1D, purple). This indicates that there is still material pinched between the beads when cells are treated with LatA. It might contain membrane, polysaccharide chains such as glycocalyx, proteins that link the membrane to the actin cortex such as ezrin/radixin/moesin, and other cytoskeletal components such as septins or intermediate filaments ^2,21^.

In conclusion, pinching the cortex with a pair of magnetic beads provide an accurate measure of the cell cortex thickness in live cells. We evidence that dendritic cells possess a thin and stiff cortical layer mostly composed of actin filaments, with properties comparable to the reported values in other cell types.

### The actin cortex thickness displays large non-periodic local instabilities

We next ask whether the cell cortex has a constant thickness, or whether thickness fluctuates in time. To address this question, we use time-resolved measures for single live cells comparing different bead configurations (Fig. 2A). Our measurement is extremely steady for beads outside the cells (control), and shows moderate fluctuations for inside beads, compatible with cell internal activity ^22^. In contrast, we observe large and fast fluctuations (several hundred nanometers in a few tens of second) for bead pairs pinching the cortex of a live cell. Most of these fluctuations are lost when cells are treated with LatA (Fig. 2A, quantified in Fig 2B, see details in Fig S2A), showing that they are driven by the activity of the actin network. Internalized beads do not change the thickness nor the fluctuations of the cortex, as these measurements do not vary with the number of beads ingested by the cells (Fig. S2D). No periodicity is observed in the fluctuations of cortex thickness, as shown by the absence of peaks in the auto-correlation function (Fig. S2F) and of any emerging frequency in the Fourier analysis (Fig S2E). Characteristic time scales for fluctuations can nonetheless be extracted, the median of their distribution being 20s. In some rare cases (n=4 for control cells) the cortical layer is pinched at two different locations by two independent bead pairs (Fig S2G). No correlation between the signals of the two bead pairs is observed (Fig. S2H), showing that thickness fluctuations are local rather than resulting from a global cell contraction. Cumulative distribution (Fig. S2B) confirms the trend of large fluctuations in cortex thickness, these fluctuations being strongly diminished when analyzing bead pairs inside cells or at the cortical layer of actin-depolymerized cells. Remarkably, this analysis further reveals an asymmetry in these active cortex fluctuations: fluctuations associated with a thickness increase are larger than fluctuations associated with a thickness decrease (Fig. S2C). The existence of such asymmetry implies that fluctuations that increase the thickness can be considered as ‘peaks’, reflecting transient augmentation in the thickness of the cortical layer. We can thus extract a frequency by counting the number of peaks above a relevant threshold (see Sup Mat). Control cell cortices exhibit on average 0.86 peaks per minute while this number drops to 0.19 for cells treated with LatA. Altogether these results show that the cortex is not a static structure with a constant thickness but is, on the contrary, a very dynamic object with large fluctuations in the direction perpendicular to the membrane. These fluctuations nevertheless remain below the resolution of classical imaging techniques, explaining why they had never been observed before. These measures therefore reveal a novel picture of the actin cortex as an unstable active layer that displays non-periodical events of thinning and thickening.

**Figure 2.**
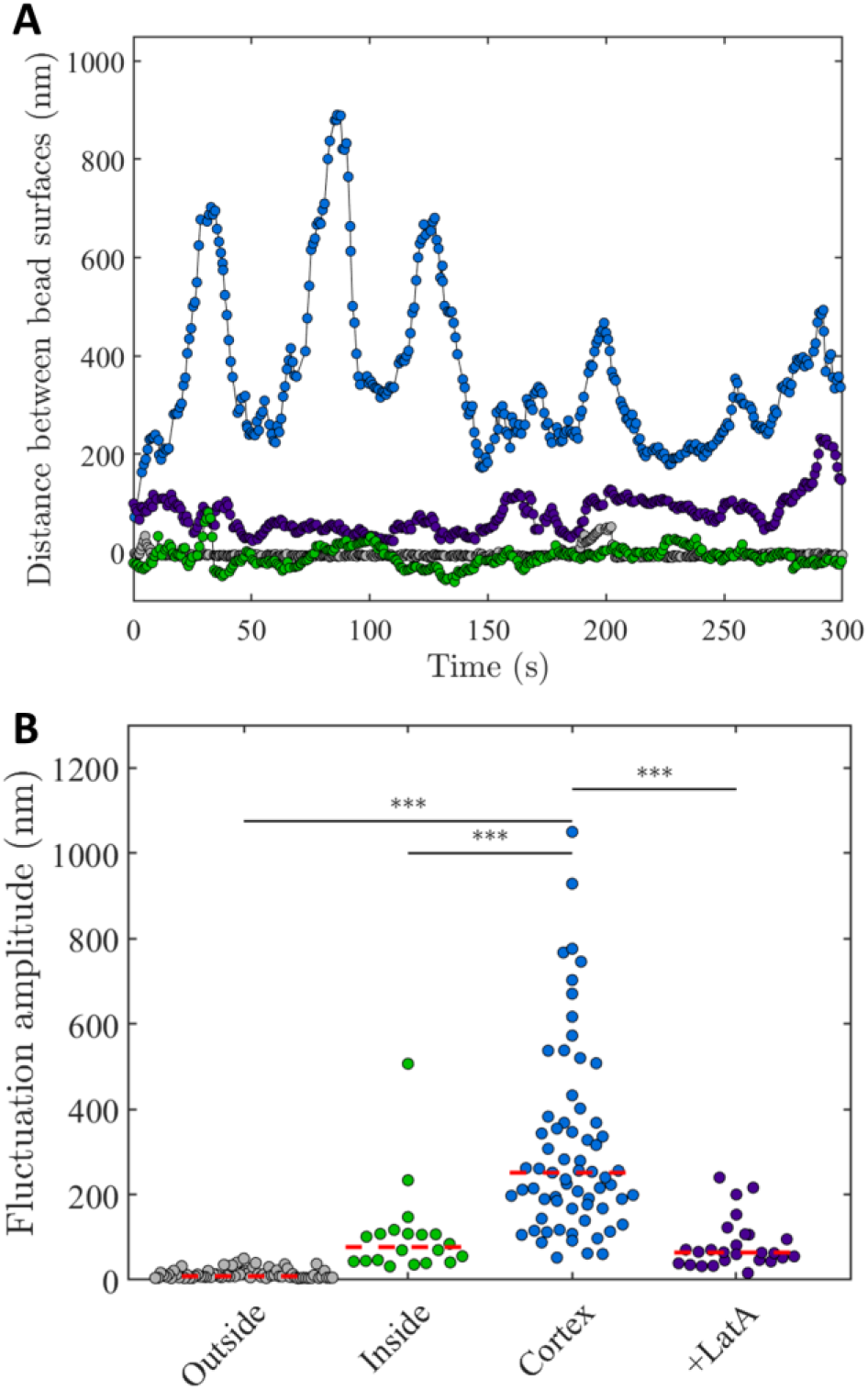
Cortex thickness fluctuates in time. A – Typical temporal evolution of the cortex thickness of control cells (blue) and LatA treated cells (purple) compared to the temporal evolution of the distance between beads surfaces inside (green) or outside (grey) cells. Acquisition rate is 1.25 Hz. Large fluctuations are visible in the measured cortex thickness of control cells (blue) but not in the other signals. B – Amplitude of the cortex thickness fluctuations in control cells (blue, n = 67, N = 10), compared to the fluctuations of the distance between bead surfaces in LatA treated cells (purple, n = 26, N = 5) and pairs of beads inside (green, n = 20, N = 13) or outside (grey, n = 69, N = 2) cells.

### Cortex fluctuations depend on actin polymerization and actomyosin contractility

While actin cortex thickness is mostly regulated by structural properties such as filament length ^10^, fast fluctuations more likely rely on active processes such as actin assembly and contractility. We thus investigate the role of actin nucleators, Arp2/3 and formin, and of the Myosin II motors in this process. We treat the cells with small inhibitors, after they had ingested the beads, as bead ingestion requires an active actin cytoskeleton.

Confocal imaging of Lifeact-GFP-expressing cells after drug treatment shows the expected effects of inhibition of actin nucleators on surface ruffles, formin inhibition having the most pronounced effect (Fig. 3A, S3F). Actin nucleation impairment from formin or Arp2/3 inhibition leads to a moderate but significant reduction of cortex thickness and to a strong decrease of the amplitude of cortex thickness fluctuations (Fig 3 C-D). This decrease is also visible in the cumulative distribution of thickness above the median (Fig S3A). In parallel with the decrease in amplitude, the number of peaks (larger than 100 nm) observed per minute dropped to almost half of the control value (Fig 3E). This tendency is more pronounced for the largest peaks (larger than 600 nm) with a 3-fold reduction for Arp2/3 and formin inhibition (Fig. 3F). Importantly, formin inhibition, which almost completely abolished membrane ruffles visible on microscopy images, had a more limited effect on cortical thickness and fluctuations. Altogether, the results on small-molecule inhibitors suggest that the sub-micron fluctuations in cortical thickness observed here do not result from the large surface ruffles, but rather correspond to fluctuations in the thickness of the cortical layer itself.

**Figure 3.**
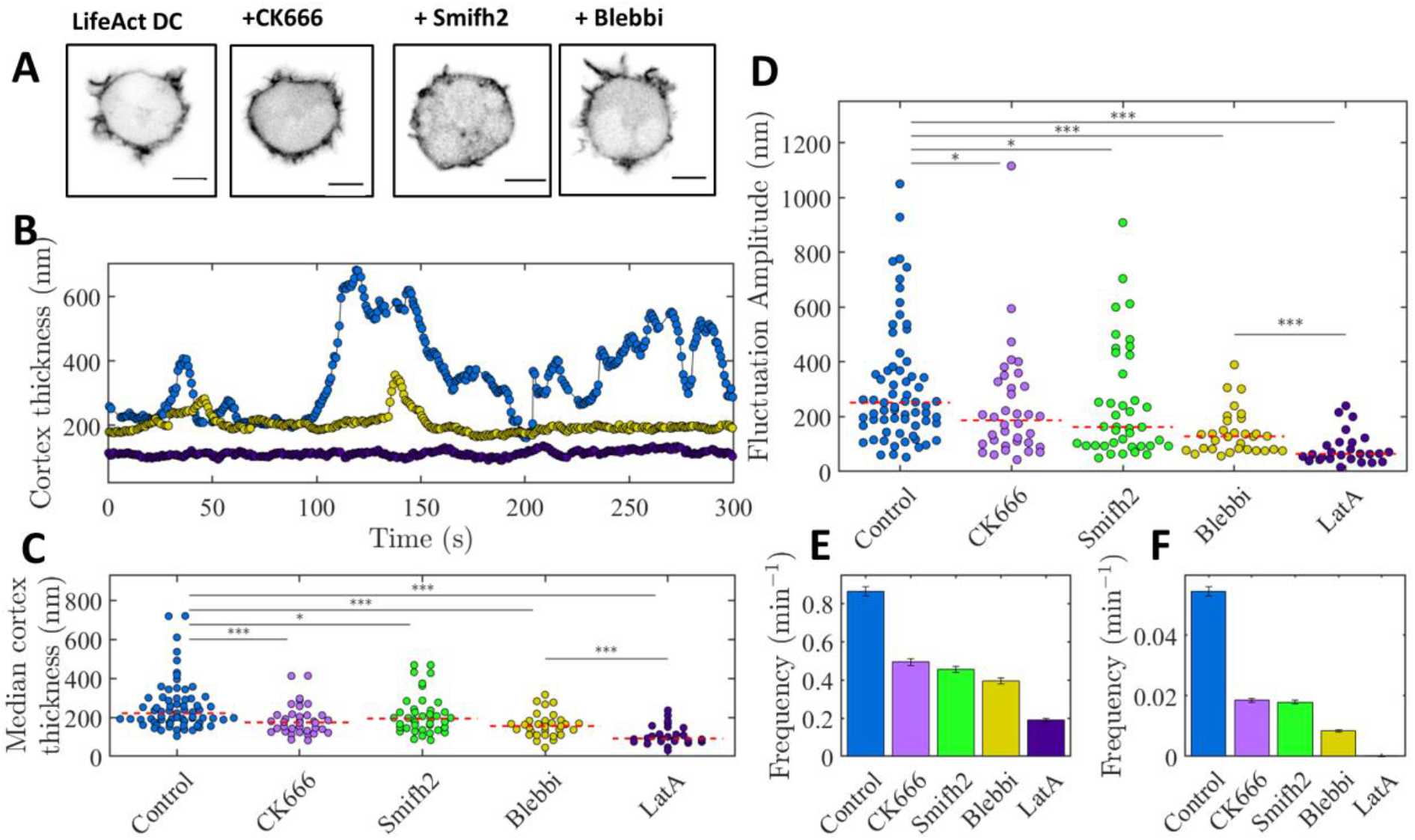
Inhibition of Myosin II affects cortex thickness and fluctuations more drastically than inhibition of nucleators. A – Confocal imaging of actin in live lifeact dendritic cells treated with DMSO, 50μM CK666, 12,5 μM SMIFH2 and 50μM blebbistatin (left to right). After treatment by CK666 protrusions appear to be sharper; Smifh2 treatment strongly reduces the number of protrusions; blebbistatin treatment slightly affect protrusions morphology. B – Typical temporal evolution of the cortex thickness in control cells (blue), blebbistatin (yellow) and Latrunculin A (purple) treated cells. C – Median cortex thickness for control cells (blue, n= 67, N = 10), CK666 (light purple, n = 36, N = 4), SMIFH2 (green n = 40, N = 5), Blebbistatin (yellow, n = 31, N = 5) and Latrunculin A (dark purple, n = 26, N = 5) treated cells. Control and LatA data are the same as in Fig1. D – Amplitude of the cortex thickness fluctuations for control and treated cells (same conditions as in C). Myosin II inhibition has the strongest effect on fluctuations, after LatA treatment, but does not affect the morphology of the cell protrusions (A) leading to the conclusion that the measured fluctuations are not the signature of protrusion but rather fluctuations of the thickness of the underlying cortex. Control and LatA data are the same as in Fig2. E – Frequency of peaks above 100 nm for control and treated cells. The reduced number of peak events in blebbistatin treated cortices as well as with Arp2/3 and SMIFH2 inhibition is in agreement with the trend on fluctuation amplitude shown in D. F – Frequency of peaks larger than 600nm for control and treated cells.

Inhibition of myosin II motors using blebbistatin has, surprisingly, a stronger effect than inhibition of actin nucleators. Cortical thickness is decreased by about one third and the amplitude of fluctuations divided by two (Fig. 3B-D), giving a cumulative probability of thickness variation close to the one of actin-depleted cells (Fig. S3A). The frequency of actin-dependent protrusions is the lowest of all inhibition conditions with 0.39 protrusions per minute (Fig.3E), and a 6-fold decrease in the frequency of peaks larger than 600nm (Fig 3F). As blebbistatin is photo-toxic ^23^, we used the ROCK inhibitor Y27632 (Y27) to image the cells with a reduced motor activity. We observed no visible effect on the ruffling activity of the dendritic cell membrane as compared to non-treated cells (Fig. S3E, F). However, this small molecule reproduced the effect of blebbistatin on cortex thickness and associated fluctuations (Fig S3D). We conclude that large fluctuations in the thickness of the actin cortex are strongly dependent on myosin II activity.

### A minimal physical description of the cortex recapitulating the effect of Myosin II on thickness fluctuations

Initial theoretical analysis of the actin cortex proposed that its thickness results from a balance between nucleation at the plasma membrane and bulk disassembly ^24^. A more recent analysis introduced the effect of modulation of filaments length ^10^. But none of these studies accounts for our observation of active sub-micron fluctuations of cortical thickness caused by Myosin II activity. We thus turn to an extension of the minimal description in (24) that accounts for stress anisotropies ^25^. Here, the cortex is treated as an active viscous gel, with a constant influx of material at the membrane (representing polymerization) and homogenous disassembly. The contractile property of the cortex is captured as an active stress that can be different in the directions tangential and perpendicular to the membrane, due to the alignment of actin or myosin filaments (Fig. 4A). When the anisotropy in the contraction is low, a stable profile of actin density forming a compressed layer of constant thickness near the membrane emerges.

However, when the anisotropy in the active stress exceeds a threshold so that the tangential stress is strong enough and stronger than the perpendicular one, the density profile becomes unstable. The cortex contracts laterally in an inhomogeneous manner leading to local densification of actin. This induces the formation of peaks growing perpendicularly to the cortex plane. These peaks are not stable: they slide laterally and merge with each other while new ones appear (fig 4B). The dynamical behavior of the fluctuations does not settle into a periodic state and is on the contrary, aperiodic and chaotic, which matches our experimental observations. The amplitude of the fluctuations is of the same order of magnitude as the cortex thickness and the frequency of those fluctuations is given by the characteristic depolymerization time of the cortex (~seconds), which also matches our experimental measures. The thickness fluctuations we observed in our experiments are thus a generic feature of actin networks assembling on a surface and exhibiting anisotropic contractility, which points to a major effect of contractility on cortex thickness stability.

**Figure 4.**
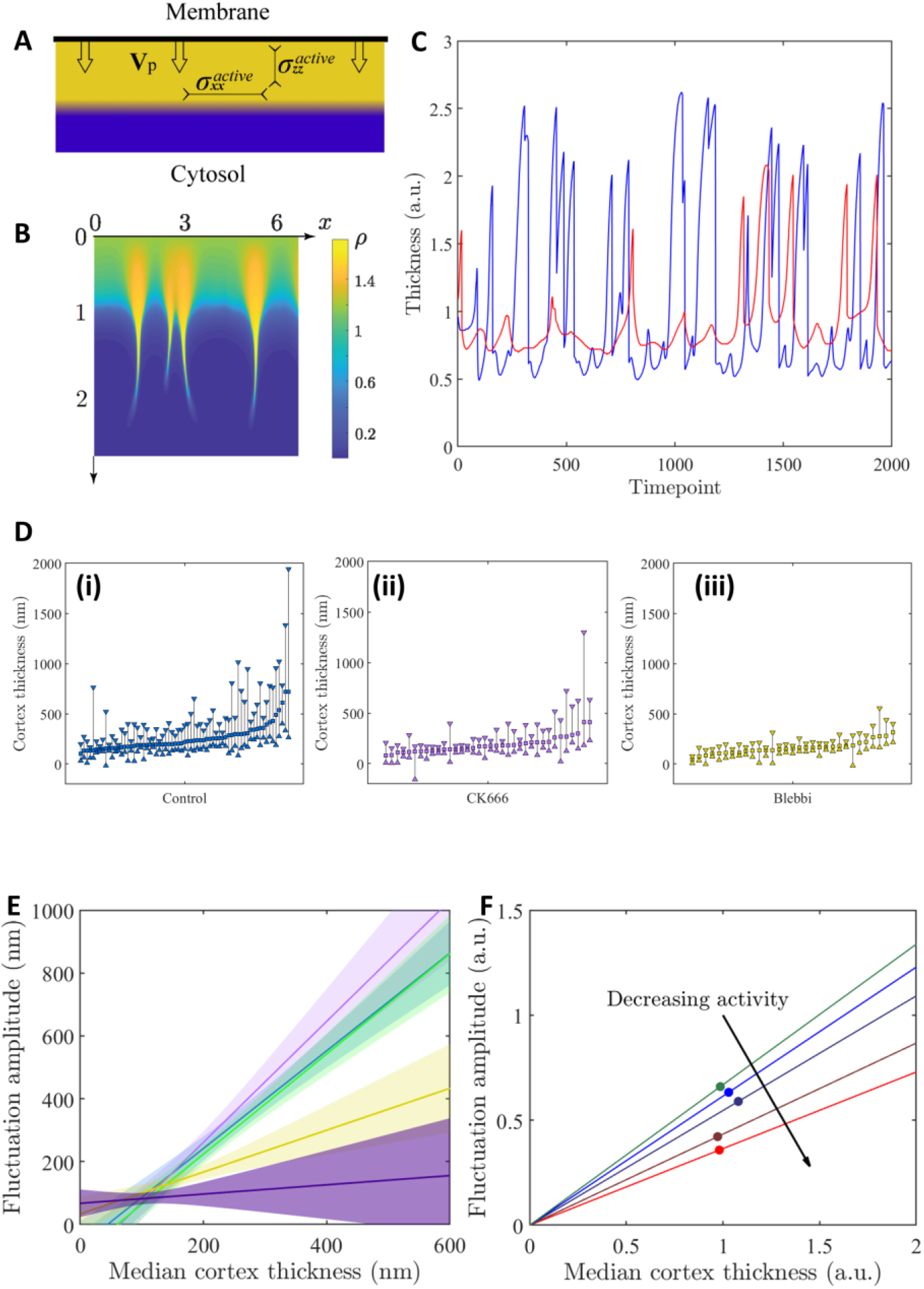
A minimal theoretical description of the cortex captures the role of contractility in thickness fluctuations. A – Illustration of the physical description of the cortex introducing the polymerization from the surface and the active stresses in the directions tangential (σ_xx_) and perpendicular (σ_yy_) to the membrane. B – Snapshot of a numerical simulation using the cortex theoretical description illustrated in A. The cortex locally deform into protruding peaks due to the stress anisotropies in the actomyosin material. C – Temporal evolution of the cortex thickness at one arbitrary coordinate from a simulation at low contractile activity (a = 7, red) and at a higher contractile activity (a = 7.75, blue). D – Representation of the cortex median thickness and fluctuation amplitude from experiments on control cells (i), CK666 (ii) and blebbistatin treated cells (iii). Cells are sorted in ascending value of the median thickness (square). The length of the vertical line between the thickness first decile (upward triangle) and last decile (downward triangle) represents the fluctuation amplitude. The thickness fluctuations appear larger for larger cortex thicknesses for control cells but not for blebbistatin treated cells. E – Slopes from linear regression of fluctuation amplitudes as a function of cortex thickness from the expriments for control cells (blue), and cells treated with SMIFH2 (green-overlapping control), CK666 (light purple), blebbistatin (yellow) and LatA (dark purple). The colored area of each curve represents half the 95% confidence interval. The correlation of fluctuation amplitude with cortex thickness is markedly stronger for control and nucleator inhibition compared to Myosin II inhibition. F – Slopes for the theoretical relation between thickness and fluctuation amplitude calculated from simulations at varying activity (dots).

To further analyze the analogy between our experimental results and the results of the theoretical analysis, we concentrate on the correlation between the cortical layer thickness and its fluctuation amplitude. In control cells, the amplitude of fluctuations in each single cell is strongly correlated with the thickness of the cortex (Fig 4D, (i)). Although both cortical thickness and its fluctuation amplitude are reduced by the inhibition of actin nucleators (Fig. 4C, (ii) & S4B, (i)), this does not affect the correlation between the two (high correlation coefficient, low p value, and slopes of the same order as for control cells, between 1.5 and 2, Fig S4A-B). On the contrary, Myosin II inhibition, which also reduces both the thickness of the cortex and the amplitude of fluctuations, has a strong effect on the correlation between the two parameters, with a lower correlation coefficient and a slope of only 0,67 (Fig. 4D,(iii)). In the case of LatA, the correlation completely disappears (p=0.55). Coming back to the simulations, we vary the active stress, and measure the cortex median thickness and its fluctuation amplitude in the same way as in the experiments (see fig 4F). As only one length scale in present in the theoretical analysis, changing the resting size of the cortex by modifying the polymerization speed will affect in the same way both the cortex median thickness and the amplitude fluctuation. Thus, in both our experiments and theory, the contractile nature of the cortex controls the correlation between cortex thickness and its fluctuation amplitude. This finding suggests that the mechanism captured by the theory can indeed explain the fluctuations in cortex thickness observed in live cells.

A large number of cell types display patterns of activity in the cell cortex, in the form of polymerization waves, global contraction or unorganized flares of activity ^26,27^. These patterns are explained by a dual mechanism of activation and inhibition in regulatory pathways, in interplay with the actin cytoskeleton ^27–29^. Myosin-dependent waves of Rho GTPase activity have been observed in adherent cells ^30^ and pulsatile patterns in embryos ^31^. Cortical flows could also entrain some larger structures embedded in the cortex and result in myosin-dependent local increase in the cortical thickness. Although we cannot rule out that these mechanisms contribute to the fluctuations of cortical thickness, the intrinsic dynamics of an active gel layer is sufficient to capture most of the characteristics of the fluctuations that we observed.

## Discussion

Collectively, our data propose a fundamentally new picture of the cell cortex in live cells as a fluctuating entity whose thickness varies on a time scale of tens of seconds with spatial amplitude of hundreds of nanometers. This picture emerges through the new method presented here that provides a time-resolved measurement of the cortex thickness of live cells with an unprecedented spatial accuracy of a few tens of nanometers. Strikingly, we found the amplitude of thickness fluctuations to be comparable to the median value of the thickness. These results are not specific to primary mouse dendritic cells, as the cortex of Dictyostelium Discoideum displays a similar behavior (see Sup Mat and Fig S7). Fluctuations mostly result from the contractile activity of myosin II motors, which produces instabilities in the cortex, inducing large shifts perpendicular to the plasma membrane. They are intrinsically different from ruffles commonly observed by fluorescence microscopy. As our measure is local, it does not distinguish whether these shifts are due to bumps or wrinkles, but depicts the cortex as a dynamically embossed structure.

What are the putative functions of these cortical fluctuations that cells could modulate through the regulation of Myosin II? One possible role could be to dynamically generate a rough plasma membrane surface that would augment the effective area of the cell surface and regulate membrane tension. Pulling membrane tethers from adherent fibroblasts revealed the existence of membrane reservoirs that depend on actin filaments ^32^. Gauthier, Masters and Sheetz ^33^ proposed that these reservoirs come from sub-micron membrane wrinkles in the plasma membrane. The fluctuations in cortex thickness here reported could provide such a reservoir. Another possible function of a wrinkled or bumped cortex could be to provide dents on the outer surface of the cell to increase its friction with the substrate. Non-adherent cells can migrate faster than adherent cells, but the way forces are transmitted to the substrate in this migration mode is still mysterious. Possessing a dynamically rough surface could help dendritic cells generate propulsion forces in low adhesion environments. Finally, the myosin-produced instabilities we evidenced give rise to fluctuations in thickness close to its median value. When these instability become too strong, the cortex could rupture, which could induce either local detachment of the plasma membrane called blebs, or even lead to large scale cell polarization ^19,31,34^.

Increasing the magnetic force between the beads provides a measurement of the cortex mechanical properties. A complete study of these properties is beyond the scope of the present paper, but we are able to provide an estimate of 7 kPa for the stiffness of the cortex (see Sup Mat). This estimation of the elastic modulus of the cell cortex is similar to measurements obtained with atomic force microscopy using tips that are considered to probe the mechanics of the cortex itself ^35^. Further studies are necessary to accurately model the contact mechanics between the beads and the cortical layer and extract a complete picture of the cortex mechanics. The precise determination of both cortex thickness and material properties is a promising avenue to elucidate the quantitative contribution of the cortex mechanics to the global deformation of the cell in response to external mechanical stresses. Another noteworthy observation is that disassembly of F-actin did not completely abolish cortex thickness, leaving a compressible layer. The estimation of the elastic modulus of the remaining layer after LatA treatment provided a value of 18 kPa (see Sup Mat), which is much stiffer than the control cortex. This stiffness may evoke strongly cohesive networks of polymers such as the intermediate filaments Vimentin, recently reported to form intertwined networks with actin filaments in the cortex of mitotic cells ^21,36^. We anticipate that in depth investigation of the link between the cell cortex mechanics, its components and external cues will be feasible by the cortex pinching method outlined here.

In conclusion, we believe that our observations not only provide measures of the physical properties of the cell cortex with unprecedented accuracy, but also draw a novel picture for this subcellular entity as an active polymorphic layer dynamically thinned and thickened on a submicron scale as a result of actomyosin contractility. This could explain the propensity of the actin cortex to break upon activation of contractility, a phenomenon used by cells to polarize and move.

## Acknowledgements

The authors wish to thanks Paula Belska for preliminary experiments on the magnetic pincher, Raphael Voituriez for useful discussions, Xavier Benoit Gonin and Olivier Brouard for technical support on the setup Aastha Mathur, Mathieu Deygas, Hélène Moreau, Doriane Sanseau, Graciela Delgado, Zahraa Alraies for additional cell culture, Nishit Srivastava for his help on Dictyostelium Discoideum culture and handling.

## Author contributions

O.d.R, M.P. and J.H. designed the research; V.L., OdR and J.H developed the cortex pinching experiments; V.L. carried out all the cortex pinching experiments and analyzed the resulting data; L.B. and P.J.S. performed differentiation and culture of dendritic cells under the supervision of P.V.; J.P. carried out all confocal imaging experiments; M.S.R. performed electronic microscopy experiments; V.L. performed live cells spinning disk imaging; V.L., L.B. and P.J.S carried out and analyzed immunofluorescence experiments; K.K. and N.L. developed the theoretical analysis; N.L. performed and analyzed cortex simulations; V.L, O.d.R., M.P., and J.H. interpreted experimental data with the contribution of P.V. and A.M.L; V.L., O.d.R., M.P. and J.H. wrote the manuscript.

## References

1. Pollard, T. D. & Cooper, J. Actin, a central player in cell shape and movement. Science (80-.). 326, 1208–1212 (2009).

2. Chugh, P. & Paluch, E. K. The actin cortex at a glance. J. Cell Sci. 131, jcs186254 (2018).

3. Salbreux, G., Charras, G. & Paluch, E. Actin cortex mechanics and cellular morphogenesis. Trends Cell Biol. 22, 536–545 (2012).

4. Diz-Muñoz, A., Weiner, O. D. & Fletcher, D. A. In pursuit of the mechanics that shape cell surfaces. Nat. Phys. 14, 648–652 (2018).

5. Fritzsche, M., Erlenkämper, C., Moeendarbary, E., Charras, G. & Kruse, K. Actin kinetics shapes cortical network structure and mechanics. Sci. Adv. 2, 1–13 (2016).

6. Bovellan, M. et al. Cellular control of cortical actin nucleation. Curr. Biol. (2014). doi:10.1016/j.cub.2014.05.069

7. Biro, M. et al. Cell cortex composition and homeostasis resolved by integrating proteomics and quantitative imaging. Cytoskeleton 70, 741–754 (2013).

8. Svitkina, T. M. Actin Cell Cortex: Structure and Molecular Organization. Trends Cell Biol. 1–10 (2020). doi:10.1016/j.tcb.2020.03.005

9. Clark, A. G., Dierkes, K. & Paluch, E. K. Monitoring Actin Cortex Thickness in Live Cells. Biophysj 105, 570–580 (2013).

10. Chugh, P., Clark, A. G., Smith, M. B. & Davide, A. D. Actin cortex architecture regulates cell surface tension. Nat. Cell Biol. 19, (2017).

11. Clausen, M. P., Colin-York, H., Schneider, F., Eggeling, C. & Fritzsche, M. Dissecting the actin cortex density and membrane-cortex distance in living cells by super-resolution microscopy. J. Phys. D. Appl. Phys. 50, (2017).

12. Vargas-Pinto, R., Gong, H., Vahabikashi, A. & Johnson, M. The effect of the endothelial cell cortex on atomic force microscopy measurements. Biophys. J. 105, 300–309 (2013).

13. Fabry, B. et al. Scaling the microrheology of living cells. Phys. Rev. Lett. 87, 1–4 (2001).

14. Pujol, T., du Roure, O., Fermigier, M. & Heuvingh, J. Impact of branching on the elasticity of actin networks. Proc. Natl. Acad. Sci. 109, 10364–10369 (2012).

15. Belbahri, R. et al. Mechanical stiffness of reconstituted actin patches correlates tightly with endocytosis efficiency. PLoS Biol. 17, 1–19 (2019).

16. Renkawitz, J. et al. Adaptive force transmission in amoeboid cell migration. Nat. Cell Biol. 11, 1438–1443 (2009).

17. Lämmermann, T. et al. Rapid leukocyte migration by integrin-independent flowing and squeezing. Nature 453, 51–55 (2008).

18. Vargas, P. et al. Innate control of actin nucleation determines two distinct migration behaviours in dendritic cells. Nat. Cell Biol. 18, 43–53 (2015).

19. Liu, Y. J. et al. Confinement and low adhesion induce fast amoeboid migration of slow mesenchymal cells. Cell 160, 659–672 (2015).

20. Bauër, P. et al. A new method to measure mechanics and dynamic assembly of branched actin networks. Sci. Rep. (2017). doi:10.1038/s41598-017-15638-5

21. Serres, M. P. et al. F-Actin Interactome Reveals Vimentin as a Key Regulator of Actin Organization and Cell Mechanics in Mitosis. Dev. Cell 52, 210–222.e7 (2020).

22. Guo, M. et al. Probing the stochastic, motor-driven properties of the cytoplasm using force spectrum microscopy. Cell (2014). doi:10.1016/j.cell.2014.06.051

23. Mikulich, A., Kavaliauskiene, S. & Juzenas, P. Blebbistatin, a myosin inhibitor, is phototoxic to human cancer cells under exposure to blue light. Biochim. Biophys. Acta - Gen. Subj. 1820, 870–877 (2012).

24. Joanny, J. F., Kruse, K., Prost, J. & Ramaswamy, S. The actin cortex as an active wetting layer Active Matter. Guest editors: Ramin Golestarian, Sriram Ramaswamy. Eur. Phys. J. E 36, (2013).

25. Levernier, N. & Kruse, K. Spontaneous formation of chaotic protrusions in a polymerizing active gel layer. New J. Phys. 22, (2020).

26. Yang, Y. & Wu, M. Rhythmicity and waves in the cortex of single cells. Philos. Trans. R. Soc. B Biol. Sci. 373, (2018).

27. Bement, W. M. et al. Activator-inhibitor coupling between Rho signaling and actin assembly make the cell cortex an excitable medium. Nat Cell Biol. 176, 139–148 (2015).

28. Devreotes, P. N. et al. Excitable Signal Transduction Networks in Directed Cell Migration. Annu. Rev. Cell Dev. Biol. 33, 103–125 (2017).

29. Pal, D. S., Li, X., Banerjee, T., Miao, Y. & Devreotes, P. N. The excitable signal transduction networks: Movers and shapers of eukaryotic cell migration. Int. J. Dev. Biol. 63, 407–416 (2019).

30. Graessl, M. et al. An excitable Rho GTPase signaling network generates dynamic subcellular contraction patterns. J. Cell Biol. 216, 4271–4285 (2017).

31. Nishikawa, M., Naganathan, S. R., Jülicher, F. & Grill, S. W. Controlling contractile instabilities in the actomyosin cortex. Elife 6, 1–21 (2017).

32. Raucher, D. & Sheetz, M. P. Membrane expansion increases endocytosis rate during mitosis. J. Cell Biol. 144, 497–506 (1999).

33. Gauthier, N. C., Masters, T. A. & Sheetz, M. P. Mechanical feedback between membrane tension and dynamics. Trends Cell Biol. 22, 527–535 (2012).

34. Ruprecht, V. et al. Cortical contractility triggers a stochastic switch to fast amoeboid cell motility. Cell 160, 673–685 (2015).

35. Wu, P. H. et al. A comparison of methods to assess cell mechanical properties. Nat. Methods 15, (2018).

36. Duarte, S. et al. Vimentin filaments interact with the actin cortex in mitosis allowing normal cell division. Nat. Commun. 1–19 (2019). doi:10.1038/s41467-019-12029-4

