## Supplementary material for "Pinching the cortex of live cells reveals thickness instabilities caused by Myosin II motors": Methods & Supplementary Materials

**Materials.** The superparamagnetic beads (Dynabeads M-450) were purchased from Invitrogen (Carlsbad, CA, USA). DMSO was purchased from Sigma-Aldrich (St-Louis, MO, USA); SMIFH2, Blebbistatin, CK666 and LatrunculinA were purchased from Tocris Bioscience (Bristol, UK).

**Magnetic Setup.** The setup is mounted on an Axio A1 inverted microscope (Carl Zeiss, Germany) with an oil-immersion 100x objective (1.4 NA) mounted on a piezocontroled translator (Physik Instrumente, Karlsruhe, Germany). The magnetic field is generated by two coaxial coils (SBEA, Vitry, France) with mu metal core (length 40 mm, diameter 26–88 mm,

750 spires). The coils are powered by a bipolar operational power supply amplifier 6A/36V (Kepco, Flushing, NY) controlled by a data acquisition module (National Instruments, Austin, TX). The maximum field generated is 100 mT with a gradient less than 0.1 mT·mm$^{-1}$ over the sample. The chains of beads are formed with a constant field of 5 mT. Time lapse images are recorded by an Orca Flash4 CMOS camera (Hamamatsu Photonics, Hamamatsu, Japan). The setup was heated at 37°C using *the box* and *the cube* from Life Imaging Systems (Basel, Switzerland). The setup is controlled by a custom Labview interface that ensures the synchronicity between piezo position, magnetic field imposition and image acquisition.

**Cells.** Immature mouse bone-marrow-derived dendritic cells were obtained by differentiation of bone marrow precursors for 10-11 days in dendritic cell medium (IMDM, FCS (10%), Glutamine (20 mM), pen–strep (100U $ml^{-1}$) and 2-ME (50 mM) supplemented with granulocyte–macrophage colony stimulating factor (50 ng $ml^{-1}$)-containing supernatant obtained from transfected J558 cells, as previously described (*37*).

**Mice.** *LifeAct-GFP* mice were a gift from M. Sixt (IST, Austria) (*38*) and bred in Institut Curie animal facilities. In general, 6 to 10 weeks old mice were used as source for bone marrows to generate dendritic cells as described above. For animal care, the European and French National Regulation for the Protection of Vertebrate Animals used for Experimental and other Scientific Purposes (Directive 2010/63; French Decree 2013-118) were strictly followed. The present experiments, which used mouse strains displaying non- harmful phenotypes, did not require a project authorization and benefited from guidance of the Animal Welfare Body, Research Center, Institut Curie.

**Cell fluorescent imaging.** Spinning disk images were acquired on a Leica DMi8 microscope (Leica, Wetzlar, Germany) with an Orca Flash4 camera and a 100X objective. The setup was heated at 37°C using *the box* and *the cube* from Life Imaging Systems. Acquisition was done using the Metamorph software. Confocal images were acquired using a laser scanning microscope Leica TCS SP8 with a 40X oil immersion objective (1.4 NA), the HyD detector (Leica) and the LAS-AF software.

**Sample preparation.** Dynabeads M450 were washed three times in milliQ water and stored in dendritic cell medium. Cells were incubated at $5.10^5$ cells per ml, with coated Dynabeads (1:1 ratio) for 1h at 37°C in dendritic cell medium supplemented with 20 mM HEPES (Sigma). After 1 hour, 27+-4 % of the cells have ingested at least one bead. Observation was done in home-made PDMS chambers coated with 1% BSA (Sigma). Cells are observed during a maximum of 1h30 with no measured differences between the beginning and the end of the experiment.

For TEM experiments cells were plated on µ-dishes with a glass bottom containing an imprinted 50 µm cell location grid (biovalley, ref 81148). Cells were fixed in 2.5% glutaraldehyde in 0.1 M Na-Cacodylate buffer pH 7.4 for 1h, postfixed for 1h with 1% buffered osmium tetroxide, dehydrated in a graded series of ethanol solution, and then embedded in epoxy resin.

**Drugs treatment.** CK666 and Blebbistatin were used at 50µM, SMIFH2 at 12.5µM, Latrunculin A at 500nM and the same quantity of DMSO was used in control experiments. PDMS chambers were pre-incubated with drugs 30 min before cell loading. Drugs were added to cell suspension at least 30 minutes before the beginning of the experiment.

**Determination of the bead distance.** Image analysis is carried out using ImageJ (NIH, USA) and Matlab (MathWorks, MA, USA). The in-plane position of each particle is determined by a weighted average of gray levels (Fig S5A) giving an accuracy of 2 nm as was done for in vitro work (*14*, *20*). Such accuracy can be obtained because the size of the Airy disk spans a few tens of pixels and because we tune the intensity of the incoming light to use the whole range of the 16-bit depth of the camera. A large number of information bits can thus be used to find the sub-pixel location of the center of the bead.

The position of the bead in the vertical direction is measured using the diffraction pattern on several images at different height (*39*). Three images were recorded in quick succession (50 ms interval) at heights distant of 300 nm by moving the objective with a piezo controlled translator. The diffraction patterns were compared to a reference (a *depthograph*), which was regularly updated. These *depthographs* are obtained by generating Z-stacks of immobilized beads and concatenating the central pixel lines of each bead image. Individual *depthographs* from independent beads are then averaged together to create the final reference. The central pixel lines from the three images are Z position of each beads in an experiment is obtained by averaging on the three images the

A similar central pixel line is taken from a bead image on each of the three successive images and correlated onto the final *depthograph* (Fig. S5B). The horizontal distance between the centers of two beads is taken as the average of these three measures. The precision on this procedure is estimated at 45nm by cross-correlating independent *depthographs* and computing the average error. This leads to a precision of 7nm on the 3-dimensional distance between the bead centers.

To ensure the quality of the tracking in the Z-direction, data points are removed when a jump in Z (>700nm) between two time points (0.8 s) is detected, in order to avoid false measurements. Artefactual isolated points in the 3D distance curves were also removed (less than 0.05% for Z and 3D artefactual points). To ensure precision in the measurements of amplitude fluctuations of frequency of peaks, curves with too much removed data (> 50%) or that are too short (< 120 sec) were not considered for analysis (1% of total curves)

**Fluctuation and Peak Analysis.** The amplitude of fluctuations is computed as the difference between the $90^{th}$ and $10^{th}$ decile of a dataset. The asymmetry of a curve corresponds to (a-b)/(fluctuations amplitude) (Fig. S2A).

To ensure the validity of the fluctuation quantification, we study the cumulative distribution of the measured thickness for being below a certain value (Fig. S2B). The median thickness is removed from each cell and the shown curve is the mean of the cumulative distribution of each cell. The same trend of large fluctuations for the cortical thickness layer remains apparent. These fluctuations are strongly diminished when looking at a pair of beads inside the cell or at the cortical layer of actin-depolymerized cells. An asymmetry is also apparent in the cortical layer fluctuation, with larger fluctuations increasing the size of the cortical layer and smaller fluctuations decreasing its size. The upper decile of thickness is 1.7 times more distant from the median than the lower decile (asymmetry = 27 %). This asymmetry is not present with both beads inside the cells, or with actin-depleted cells (Fig. S2C).

To detect peaks, the signal is first smoothed using a Savitzky-Golay algorithm. The peaks are then detected as local maxima with a minimum prominence of 15 nm. Before computing statistics, peaks with a gap in time or in height (> 5s or >60% of prominence between two consecutive points) are removed from the data (less than 25%).
To avoid being dominated by noise, and to concentrate on actin-induced fluctuations, we considered for further analysis only the peaks above a threshold of 100 nm (Fig S2-I).

### Theoretical description of the cortex

For the theoretical analysis, we use the following physical description of the cortex as an active gel leading (*25*).We consider that the cortex is growing into the half space $z \geq 0$ at the surface $(x, y, z = 0)$ thanks to polymerisation. We note $\rho$ the density of the actin gel and $\mathrm{v} = (v_x, v_y, v_z)$ the velocity field. For the sake of simplicity, we assume invariance along the $y$-direction and $v_y = 0$. Three equations determine the temporal evolution of the cortex:

$$\partial_t \rho + \partial_x(\rho v_x) + \partial_z(\rho v_z) = -k\rho$$

$$\eta[2\,\partial_{xx}v_x + \partial_{xz}v_z + \partial_{zz}v_x] = \partial_x \Pi_x(\rho)$$

$$\eta[2\,\partial_{zz}v_z + \partial_{xz}v_x + \partial_{xx}v_z] = \partial_z \Pi_z(\rho)$$

The first equation accounts for mass conservation and the two last ones for force balance. Here, $k$ denotes the gel's disassembly rate, $\eta$ the viscosity, and $\Pi_{x,z}$ are the components of the non-viscous contribution to the total stress in the gel. The latter has two contributions: an effective hydrostatic pressure and the contractile stress generated by active processes in the gel. Both components depend on the gel density and we write (*24*, *25*):

$$\Pi_{x,z} = -a_{x,z}\rho^3 + b\rho^4$$

with $b > 0$ accounting for positive hydrostatic pressure and $a_{x,z} > 0$, which reflects contractility of the active stress component.

These equations are complemented by the boundary conditions $v_z(z=0) = v_p$ and $v_x(z = 0) = 0$, where $v_p$ denotes the polymerisation speed. See (*25*) for a discussion about more general boundary conditions including friction between the cortex and the membrane. Note that the sole length scale of this description is $l = v_p k^{-1}$. This implies that for a given value of $a_{x,z}/\eta$ and $b/\eta$, the amplitude of the fluctuations and the median thickness are proportional.

The set of previous equations generate spontaneously chaotic protrusions from the cortical layer if the active parameter $a_x$ is large enough (*25*). Figure 4 (B,C,F) is obtained by solving numerically the above equations. To this end we used a discrete Euler scheme: in each time step, we first determine the velocity field through the force balance equations, where we use Fourier decomposition along *x* and finite-differences along *z*. We then update the density. The contribution of $\partial_z(\rho v_z)$ is obtained by an up-wind finite-differences scheme in real space. To improve the stability of the scheme, we have added a small diffusion term with diffusion constant $D = 10^{-3}$ to the mass conservation equation. In all simulations, we use $\Delta x = 0.004$, $\Delta z = 0.007$, and $\Delta t = 0.0005$. Finally, we define the thickness of the cortex by the smallest $z$ value for which the actin density dropped to the half of its value at $z = 0$. Note that choosing another criterium does not affect the qualitative behaviour of our results.

### Estimation of the mechanical properties of the cell cortex

**We measure a value of 7 kPa for the elastic modulus of the dendritic cell's cortex.**

To provide an estimate of the cortex rigidity, we want to compute an elastic modulus ($E$) from the compression experiments of the cortical layer. The elastic modulus is a material property that defines how it deforms when subjected to an external force. In the case of an elastic material this modulus is the slope of the stress-strain curve, where the stress is the applied force divided by the contact surface, and the strain is the relative deformation of the material.

In our case, with $h_0$ the resting thickness of the cortex and $h$ the current thickness under force, we define the indentation $\delta = h_0 - h$ and the strain $\varepsilon = \frac{\delta}{h_0}$. In order to compute the stress, we need to know the contact area between the beads and the cortex. Due to the spherical shape of the beads, the contact area depends on $\delta$. We thus use the same geometrical consideration as in the Hertz model where the contact surface is a disc of radius $a = \sqrt{R\delta}$ and area $S = \pi a^2$ (*40*). However, the Hertz model considers the indentation of an infinite half-plane, where the deformation extends on a length scale $a$, but in this case $a > h_0$ as soon as the indentation exceeds a few nm. We thus need to consider the full layer of cortex to be under compression in order to compute an elastic modulus. To this end, we use the extension of the Hertz model for a thin slipping layer derived by Chadwick (*41*) to fit the experimental compression curves (Fig. S6A). The equation given by Chadwick for the indentation $\Delta$ of a thin layer of thickness $H_0$ resting unbounded on a substrate is:

$$F = \frac{2\pi E R \Delta^2}{3H_0} \quad (1)$$

Our situation differs from this as we have two beads indenting each side of the cortex. We thus consider a symmetric situation where two layers of thickness $\frac{h_0}{2}$ are adjacent and both indented by $\frac{\delta}{2}$. We thus obtain the equation:

$$F = \frac{\pi E R \delta^2}{3h_0} \quad (2)$$

To reduce the effect of potential non-linear elastic behavior of the cortex, we chose to fit the strain only up to 25% ($0 < \varepsilon < 0.25$, Fig. S6A). For each compression curve we obtain a value for the elastic modulus as well as a confidence interval for this value. The average elastic modulus for the dendritic cell cortex is computed with a weighted average of the modulus E, with a weight $(\frac{E}{dE})^2$ where dE is the size of the confidence interval (Fig. S6B). For control cells this average elastic modulus is 7 kPa.

**When actin is depolymerized, the remaining layer has an elastic modulus of 18 kPa**

We performed the same compression experiment on dendritic cells treated with 500 nM LatA. By using the same fitting analysis, we were able to compute an elastic modulus for the cortex layer remaining after actin depolymerization. For this layer the modulus computed using a weighted average is higher than for the complete cortex, at 18 kPa (Fig. S6C-D)

**At 5 mT the cortex is indented by 13% and the contact area is around 0.23 µm² and**

Using the values for the cortex thickness measured with a 5mT magnetic field (220 nm), the corresponding force (70pN) and the elastic modulus of the cortex (7kPa) we use the Chadwick equation (1) to compute the deformation at this level of force. We estimated this slight deformation of the cortex and underestimation of its thickness to be 33 nm, or 13% of the cortical layer thickness compared to a non-deformed layer. The low attractive force use in the experiments thus ensures a steady hold on the cortical layer without affecting excessively the accuracy of the obtained values.

We also use this estimated indentation to compute the radius of the contact surface, and its area from the Hertz model. We obtain a radius of $a = 271\ \mathrm{nm}$ and a contact surface of $S = 0.23\ \mu m^2$.

### The cortex of Dictyostelium Discoideum has dynamical properties similar to the one of dendritic cells.

In the same way as was done for mouse dendritic cells, we measure the cortex thickness and its variations in another cell type: Ax3 strain of Dictyostelium Discoideum cells. The experimental protocol is the same as for dendritic cells with 1h30 of incubation with beads. The experiment was performed on wild type cells and gave results similar to what was found for dendritic cells with a cortex thickness of 261 nm (Fig. S7A). Fluctuations however were stronger in Dictyostelium with a median of 371 nm (Fig S7B). The correlation between cortex thickness and fluctuations is also present in Dictyostelium, with a slope of 1.67, comparable to 1.55 for dendritic cells (Fig. S7C). All together these results indicate that dendritic cells and Dictyostelium cells have a similar cortex behavior. An important next step would be to confirm the role of myosin in Dictyostelium, as well as testing other cell types, especially mesenchymal cells in order to compare their cortical behavior.

Figure S1

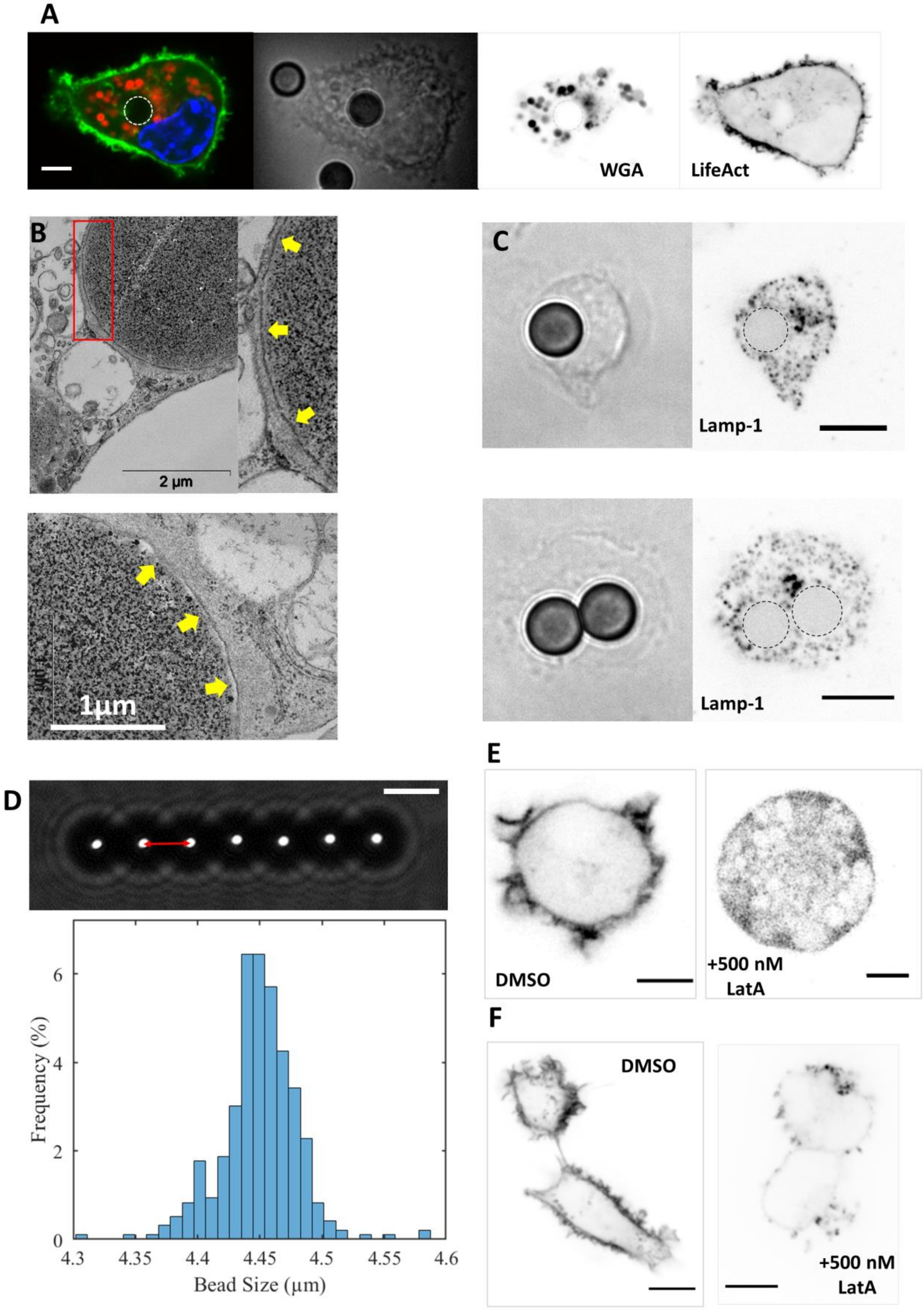
A
WGA
LifeAct
B
2 µm
1µm
C
Lamp-1
Lamp-1
D
Frequency (%)
Bead Size (µm)
0
2
4
6
4.3
4.35
4.4
4.45
4.5
4.55
4.6
E
DMSO
+500 nM LatA
F
DMSO
+500 nM LatA

*A – Spinning disk images of live dendritic cells with ingested magnetic beads. Actin is fluorescently labelled in green (lifeact), lyososomes in red (WGA) and nucleus in blue (Hoechst). WGA staining shows that beads are not in lysosomes. LifeAct shows that there is no actin shell around the bead.*

*B- Transmission electron micrographs showing a layer of membrane around beads inside a cell (yellow arrows). On the top row, right image is an enlargement of the red region in the left image.*

*C – Bright-field (left) and spinning disk (right) images of two different fixed dendritic cells with ingested magnetic beads. On the fluorescent image, lamp-1 antibody is staining lyososomal membranes. We note the absence of lysosomal signal around the beads, in agreement with (A).*

*D – Size distribution of the M450 (Dynal) magnetic beads used in the magnetic pincher experiments. This center-to-center distance, which correspond to the bead diameter, is measured on bright field images like the one on the top row where one can see the chain that beads spontaneously form in a high magnetic field. The distribution is centered on 4450 +-2 nm, with a standard deviation of 30 nm. This diameter is subtracted from the distance between the centers of the two beads that pinch the cell cortex to measure its thickness. The very low polydispersity measured here is the main source of the random error on the cortex thickness measurement.*

*E – Confocal image of actin (Lifeact) in live dendritic cells: Left, control, (DMSO), right, and 500nM Latrunculin A (right). Contrast on Latrunculin A has been increased a lot as compared to control to reveal relevant features. In the LatA case there are no more actin structures protruding from the cell and no more cortical actin is visible.*

*F – Spinning disk imaging of actin (Phalloidin) in fixed dendritic cells treated with DMSO (left) and 500nM LatrunculinA (right). Observation conditions and contrast adjustment are identical for both images.*

Figure S2

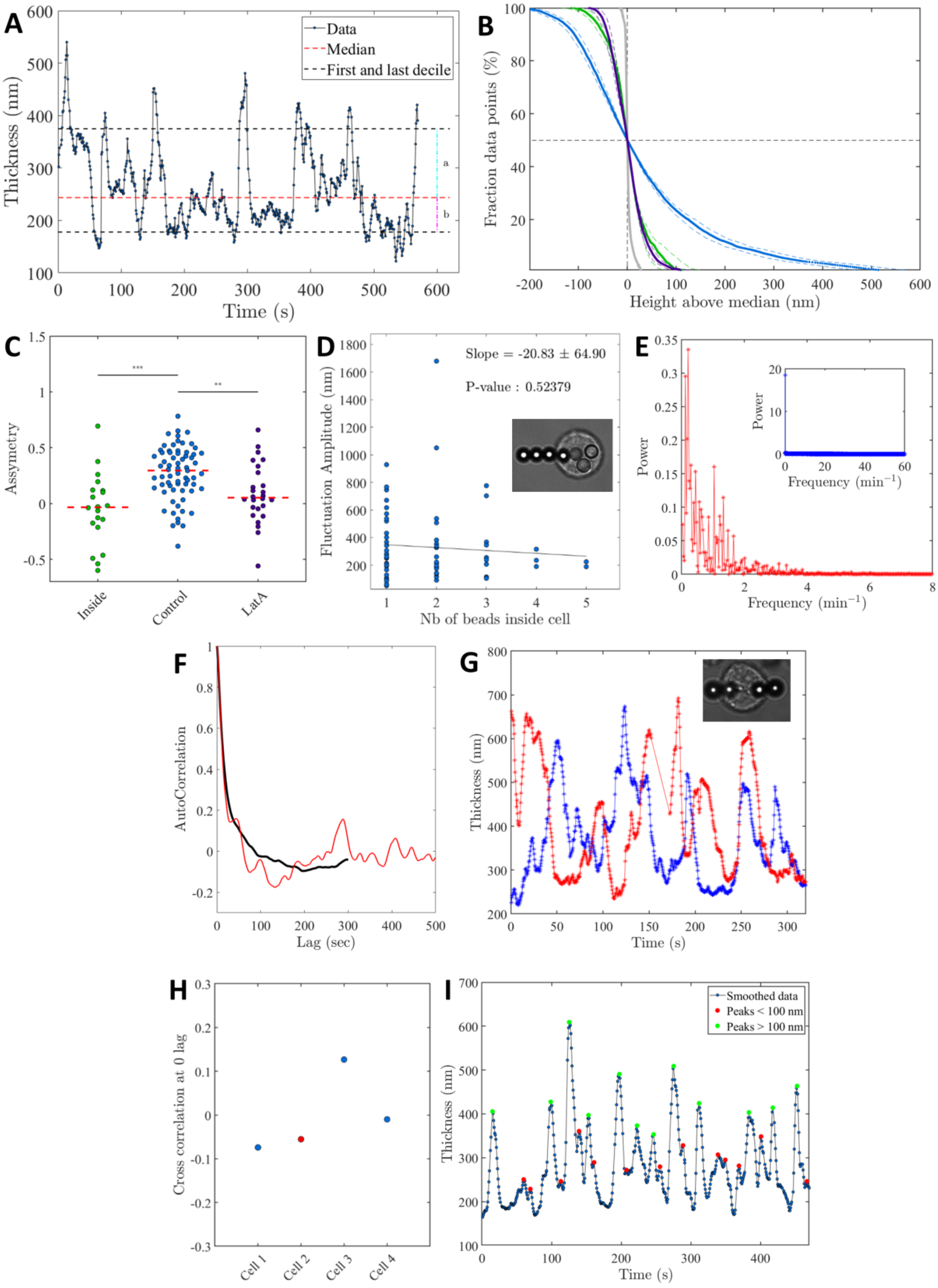

A
Thickness (nm)
Time (s)
Data
Median
First and last decile
a
b
B
Fraction data points (%)
Height above median (nm)
C
Assymetry
Inside
Control
LatA
D
Fluctuation Amplitude (nm)
Nb of beads inside cell
Slope = -20.83 ± 64.90
P-value : 0.52379
E
Power
Frequency (min⁻¹)
F
AutoCorrelation
Lag (sec)
G
Thickness (nm)
Time (s)
H
Cross correlation at 0 lag
Cell 1
Cell 2
Cell 3
Cell 4
I
Thickness (nm)
Time (s)
Smoothed data
Peaks < 100 nm
Peaks > 100 nm

*A – Typical temporal evolution of the cortex thickness for a control cell. Dotted red line indicates the median while black dotted lines correspond to the first and last decile of the data distribution. The amplitude of fluctuations is the distance between first and last decile (a+b).*

*B – Cumulative distribution of the cortex thickness centered on its median (blue). For each cell, the cumulative probability is the probability for the distance to be above the sum of its median and a certain value (in x). The cumulative distribution is averaged on all the cells considered in Fig. 2. This cumulative distribution is compared to the one of the distance between bead surfaces for LatA treated cells (purple), two beads inside (green) and outside (grey) cells.*

*C – Asymmetry of the time curve for pairs of beads inside, control and LatA treated cells. The value of asymmetry is computed as (a-b)/(a+b) as defined in A.*

*D –Amplitude of the cortex thickness fluctuations for control cells with different numbers of ingested beads. No significant effect of the number of beads inside cells is observed. Inset: example of a cell with four ingested beads.*

*E – Fourier analysis of a time curve from a 30 minute-long experiment without the 0 frequency component. Inset: zoom out of the same graph, including the 0 frequency component.*

*F – Autocorrelation of 30minute-long curve (red) and average autocorrelation curve (black) for all control cells monitored more than 5 minutes (n= 63, N= 10). The example (red) is from the same experiment as the one shown in E.*

*G – Temporal evolution of the cortex thickness pinched in two different locations on the same cell (shown in the inset).*

*H – Cross correlation at a lag of zero for the two signals obtained at different location in the same cell for four control cells. The low correlations values indicate that the thickness fluctuations do not arise from a global phenomenon. The red dot corresponds to the curves shown in S2G.*

*I – Example of a time curve with peak detection. Peaks indicated by a green (red) dots are larger (smaller) than 100nm.*

Figure S3

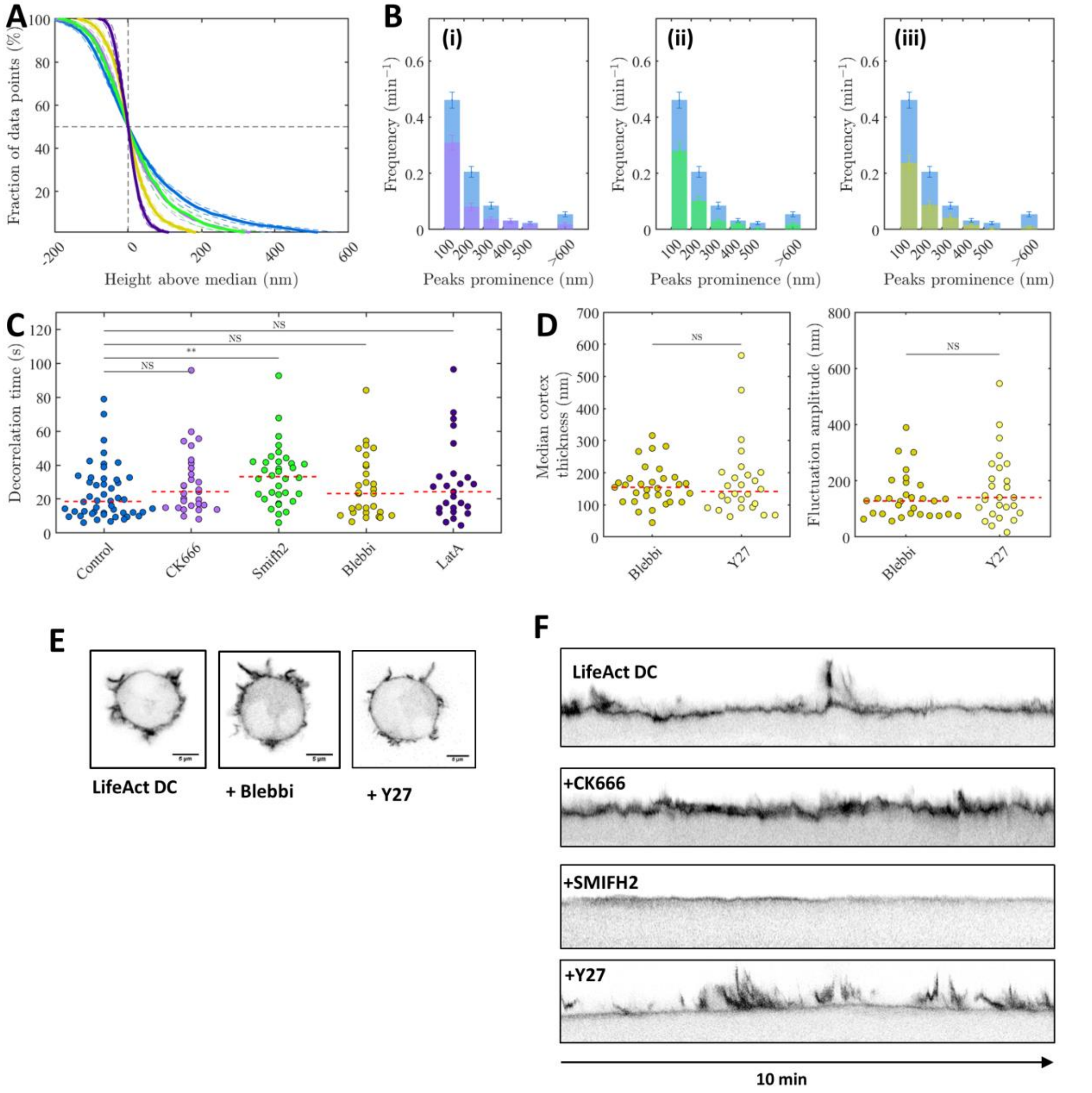


*A – Cumulative distributions of the cortex thickness above the median for control cells (blue), CK666 (light purple), SMIFH2 (green), Blebbistatin (yellow) and Latraculin A (dark purple) treated cells. Control and LatA data are the same as in FigS2.*

*B – Comparison of peak frequency as a function of peak size between control cells and CK666 (i), SMIFH2 (ii), blebbistatin (iii) treated cells.*

*C - Characteristic decorrelation time computed by fitting autocorrelation curves with a simple exponential, for control and treated cells (same experimental conditions as A). Control cells display a decorrelation time of 19s, which is increased to 30 s in the case of SMIFH2 treated cells (p=0.008) and to 24 s in the case of CK666 treated cells (non-significant, p=0.16).*

*D – Comparison of cortex thickness and fluctuation amplitude between blebbistatin and Y27 (n = 26, N = 3) treated cells. No difference between the two treatments, which inhibit myosin II motors, is observed.*

*E – Confocal imaging of actin (lifeact) in live dendritic cell treated with DMSO, blebbistatin (both same as fig3A) and Y27. Like blebbistatin, Y27 has a small effect on the number and morphology of actin protrusions.*

*F – Kymographs showing actin dynamics in the cortex for dendritic cells treated with DMSO, CK666, SMIFH2 and Y27. Kymographs were obtained from confocal imaging time-lapse (see fig3A and S3E). The absence of correspondence in the effect of inhibitors (particularly Y27 and SMIFH2) on large ruffle dynamics and submicron cortex fluctuations shows the different nature of these fluctuations.*

Figure S4

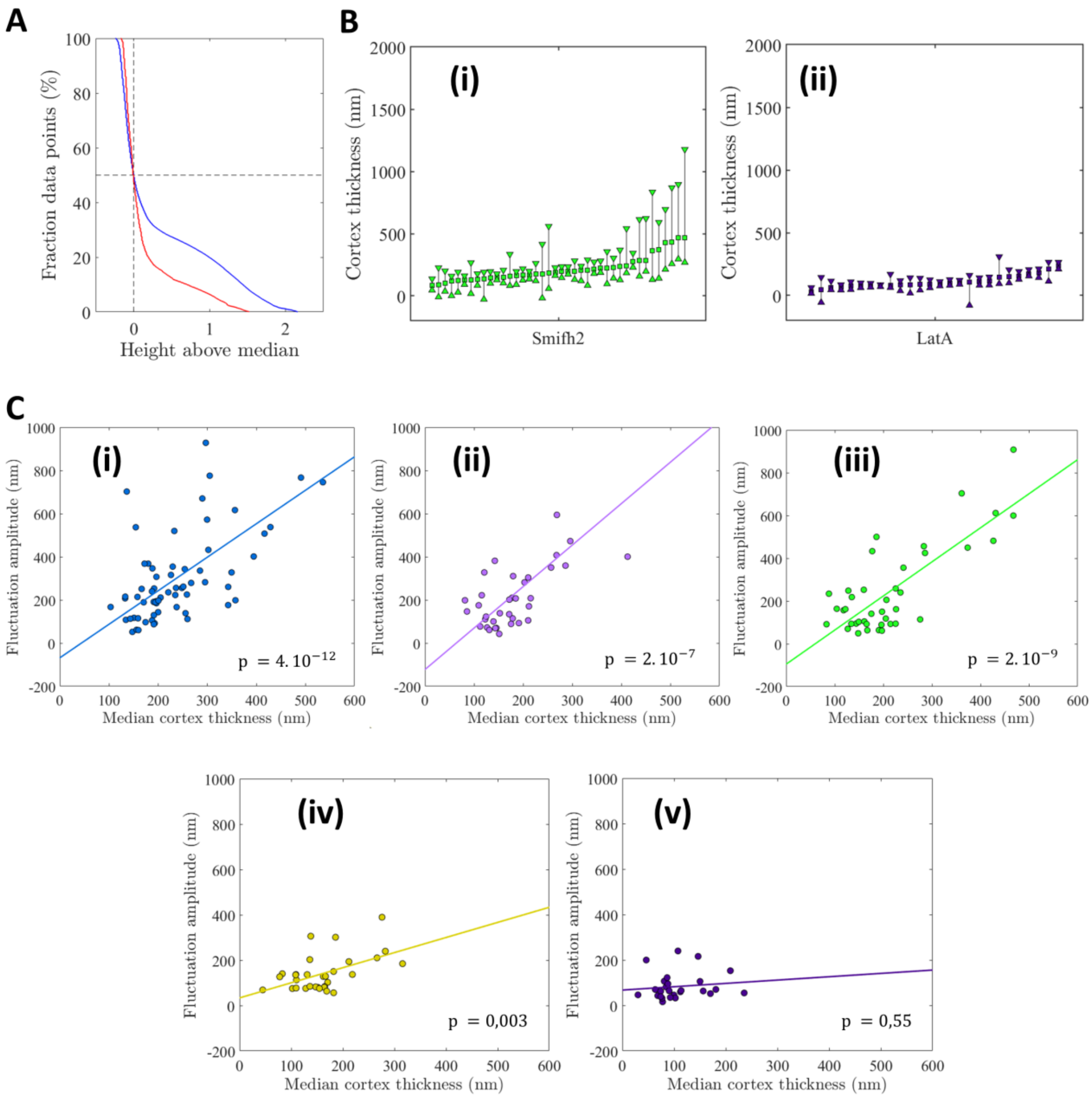
A
Fraction data points (%)
Height above median
B
(i)
Cortex thickness (nm)
Smifh2
(ii)
Cortex thickness (nm)
LatA
C
(i)
Fluctuation amplitude (nm)
Median cortex thickness (nm)
p = 4. 10^-12
(ii)
Fluctuation amplitude (nm)
Median cortex thickness (nm)
p = 2. 10^-7
(iii)
Fluctuation amplitude (nm)
Median cortex thickness (nm)
p = 2. 10^-9
(iv)
Fluctuation amplitude (nm)
Median cortex thickness (nm)
p = 0,003
(v)
Fluctuation amplitude (nm)
Median cortex thickness (nm)
p = 0,55

*A – Cumulative distribution of thickness from a simulation at low contractile activity (a = 7, red) and at a higher contractile activity (a = 7.75, blue) corresponding to Fig. 4C.*

*B– Representation of the cortex median thickness and fluctuation amplitude for Smifh2 (i), and LatA (ii) treated cells. Cells are sorted in ascending value of the median thickness (square). The length of the vertical line between the thickness first decile (upward triangle) and last decile (downward triangle) represents the fluctuation amplitude.*

*C - Linear regression for fluctuation amplitude and cortex thickness in control cells (i), and cells treated with CK666 (ii), SMIFH2 (iii), blebbistatin (iv) and LatA (v).*

Figure S5

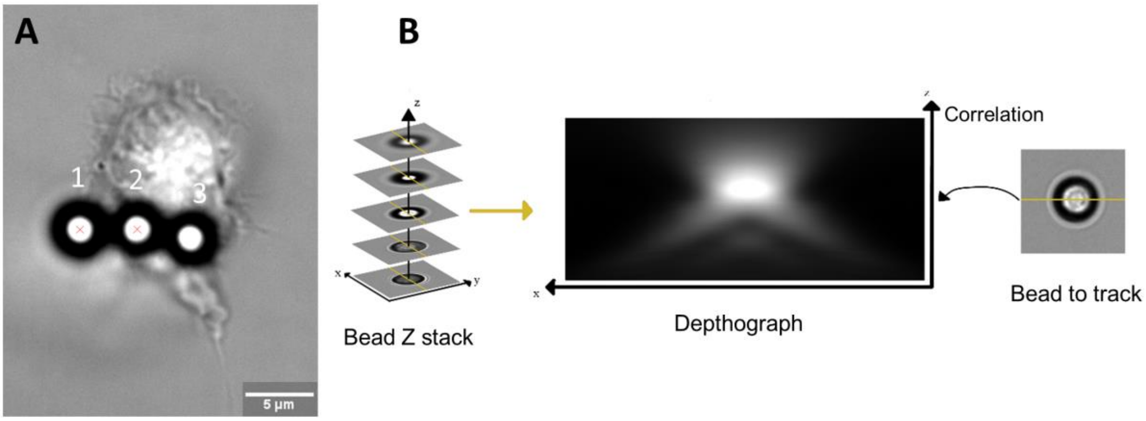


***Principle of beads tracking in 3D***

*A – Transmission image of the same cell as in fig.1B, in the plane used to track the beads. The in plane distance between the beads is computed by tracking the light spots of the beads, as indicated by the two red crosses on beads 1 and 2.*

*B – Principle of the tracking algorithm in the Z direction. Starting from a Z stack of an immobilized bead (left), a reference 'depthograph' (middle) can be created. Then, the image of a bead (right) can be correlated on the depthograph to get a Z position.*

Figure S6

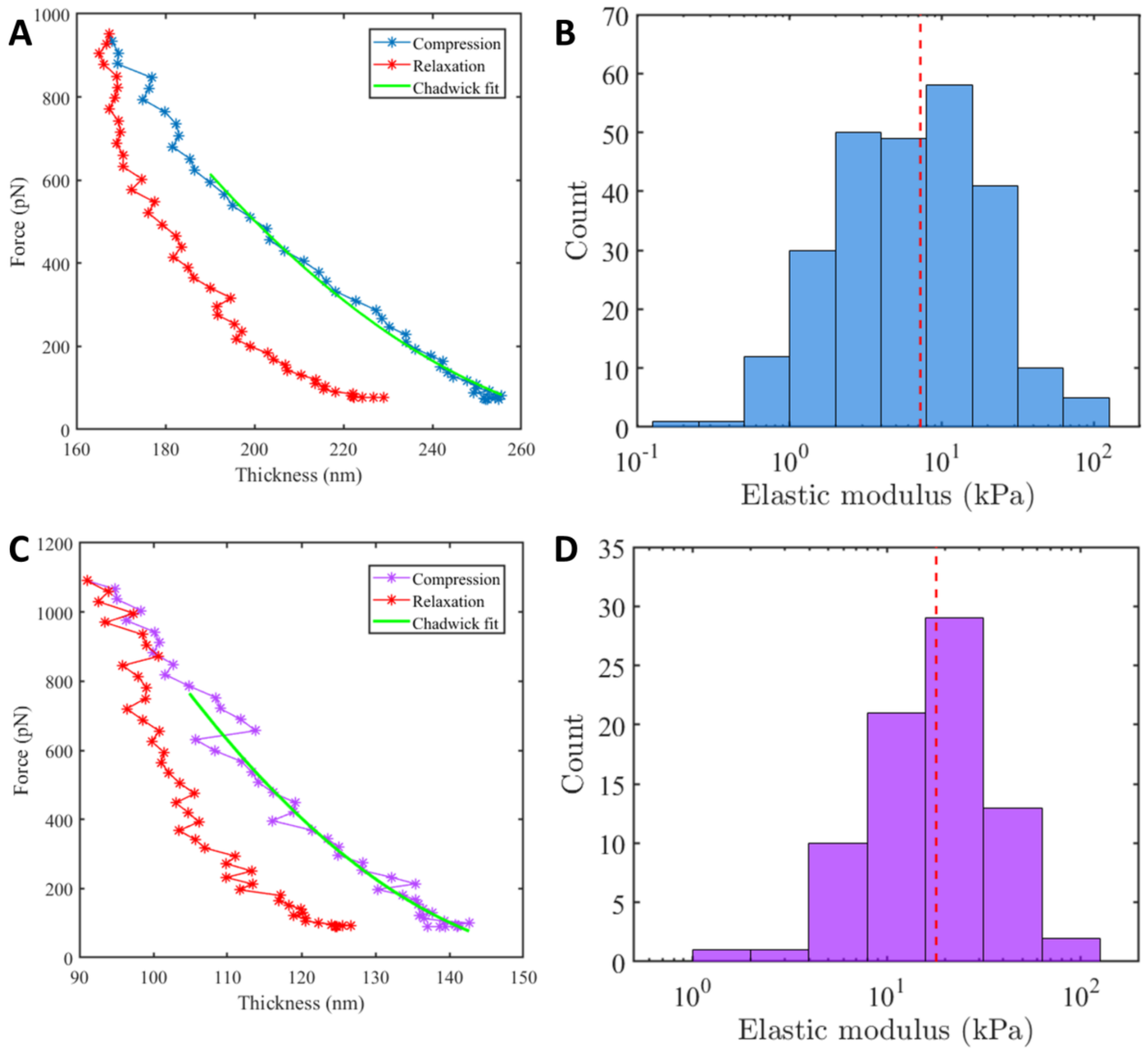


***Compression of the cell cortex gives a hint at its mechanical properties.***

*A – Example of a compression (blue) and relaxation (red) of an untreated dendritic cell. In green is a fit using Chadwick equation to fit the curve up to 25% of deformation.*

*B – Distribution of the elastic moduli obtained using Chadwick equation for control cells (n = 246 compression). The weighted average (red line) is at 6.95 kPa. The bin limits are set on a logarithmic scale.*

*C – Example of a compression (purple) and relaxation (red) of a LatA treated dendritic cell cortex. In green is a fit using Chadwick equation to fit the curve up to 25% of deformation.*

*D – Distribution of the elastic moduli obtained using Chadwick equation (n = 77 compression ). The weighted average (red line) is at 18.01 kPa. The bin limits are set on a logarithmic scale.*

Figure S7

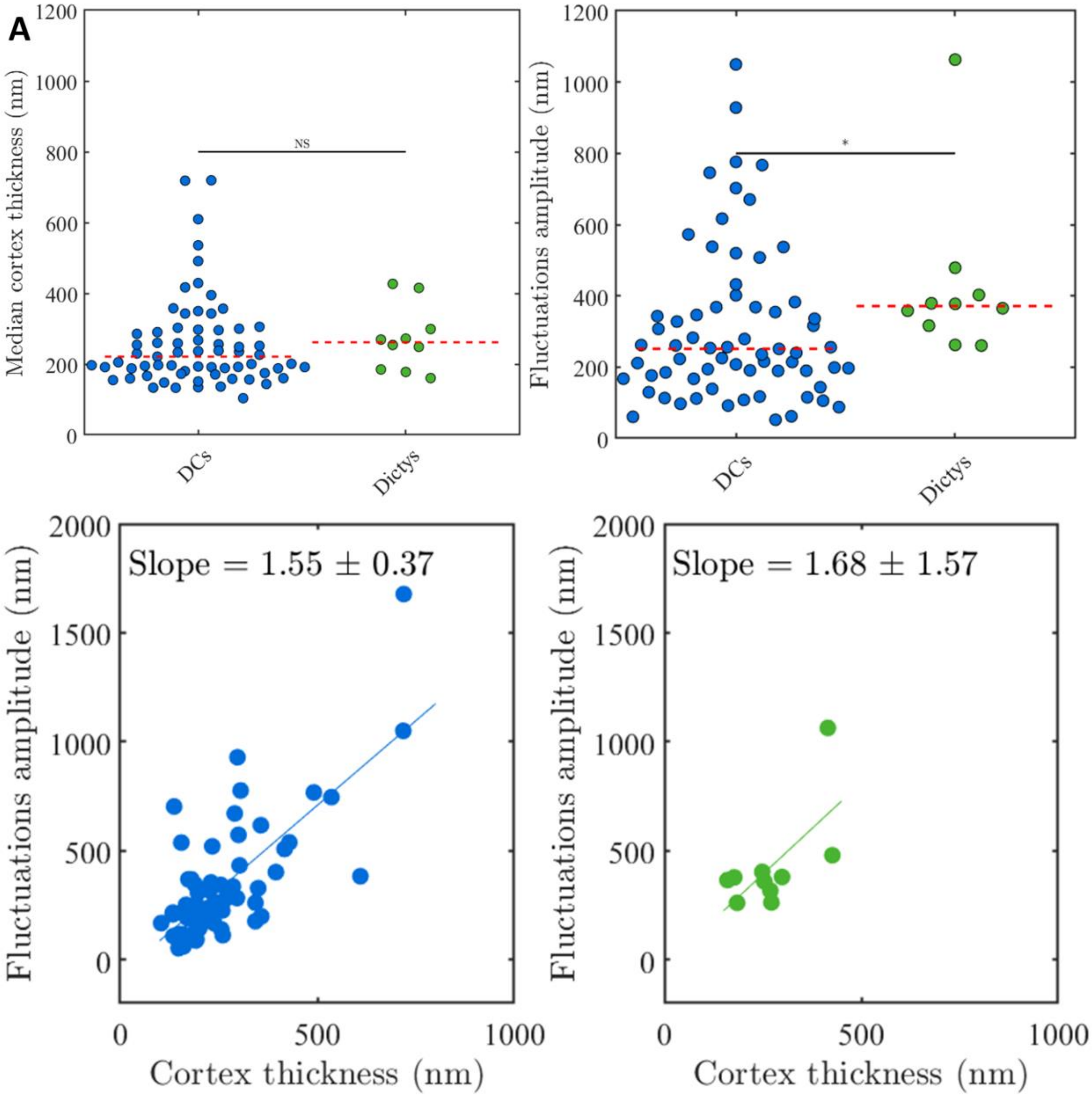


***Dendtritic cells and Dictyostelium Discoideum have similar cortex behaviors.***

*Comparison between Dentritic cells (blue, n = 67, N = 10) and Dictyostelium Ax2 (green, n = 10, N = 2) for both cortex thickness (A) and cortex thickness fluctuations amplitude (B). We also observe a similar correlation between Cortex thickness and fluctuation for dictyostelium (C).*

### References for supplementary material

37. H.-R. Thiam, P. Vargas, N. Carpi, C. L. Crespo, M. Raab, E. Terriac, M. C. King, J. Jacobelli, A. S. Alberts, T. Stradal, A.-M. Lennon-Dumenil, M. Piel, Perinuclear Arp2/3-driven actin polymerization enables nuclear deformation to facilitate cell migration through complex environments. *Nat. Commun.* (2016), doi:10.1038/ncomms10997.

38. J. Riedl, K. C. Flynn, A. Raducanu, F. Gärtner, G. Beck, M. Bösl, F. Bradke, S. Massberg, A. Aszodi, M. Sixt, R. Wedlich-Söldner, Lifeact mice for studying F-actin dynamics. *Nat. Methods*. **7**, 168–169 (2010).

39. C. Gosse, V. Croquette, Magnetic Tweezers: Micromanipulation and Force Measurement at the Molecular Level. *Biophys. J.* **82**, 3314–3329 (2002).

40. L. D. Landau, E. M. Lifshitz, Theory of Elasticity. *(Pergamon Press.* **64**, 176–177 (1960).

41. R. S. Chadwick, Axisymmetric indentation of a thin incompressible elastic layer. *SIAM J. Appl. Math.* **62**, 1520–1530 (2002).
